# The EAAT1 aspartate/glutamate transporter is dispensable for acute myeloid leukemia cell growth and response to therapy

**DOI:** 10.1101/2025.07.13.664609

**Authors:** Hernán A. Tirado, Nithya Balasundaram, Fleur Leguay, Lotfi Laaouimir, Nick van Gastel

**Affiliations:** Cellular Metabolism and Microenvironment Laboratory, de Duve Institute, UCLouvain, Brussels, Belgium; WELBIO Department, WEL Research Institute, Wavre, Belgium

**Keywords:** Aspartate, excitatory amino acid transporter, SLC1A3, AML, chemotherapy

## Abstract

Acute myeloid leukemia (AML) is an aggressive malignancy of hematopoietic stem and progenitor cells characterized by profound metabolic dysregulation. Pyrimidine biosynthesis has emerged as a critical metabolic dependency in AML, but clinical translation has been hampered by unacceptable toxicity of current pyrimidine synthesis inhibitors. Since aspartate is an essential nutrient for pyrimidine biosynthesis, we investigated the role of aspartate import via the excitatory amino acid transporter 1 (EAAT1) in AML. We found that EAAT1 is broadly expressed across AML cell lines and patient samples, with enrichment in M4 and M5 subtypes and increasing levels following chemotherapy treatment. Pharmacological inhibition of EAAT1 impaired AML cell viability *in vitro*, but metabolomic profiling and nutrient rescue experiments showed that these effects were independent of intracellular aspartate levels. Moreover, AML cells cultured in aspartate-free medium maintained proliferation and did not become more sensitive to chemotherapy. EAAT1 inhibition in mice increased bone marrow plasma aspartate levels, confirming inhibition of cellular aspartate uptake, but did not affect growth or chemosensitivity of MLL-AF9-expressing AML cells *in vivo*. These findings suggest that AML cells possess several complementary mechanisms to support their aspartate requirements and that EAAT1 inhibition does not impair AML growth or response to chemotherapy.

## Introduction

Acute myeloid leukemia (AML) is a genetically and phenotypically heterogeneous disease that arises and evolves within the bone marrow microenvironment, where both normal and malignant hematopoiesis coexist and interact (1, 2). For decades, standard AML therapy has centered on the use of cytotoxic agents such as nucleoside analogs, particularly cytarabine, which target rapidly proliferating cells by interfering with DNA replication, often in combination with anthracyclines. Over the past years, new chemotherapy regimens adapted for older patients and targeted oral therapies against driver mutations have been added to the repertoire (3). However, resistance to therapy remains a major clinical hurdle, contributing to poor long-term survival outcomes in most patients (4). While significant efforts have focused on deciphering the genetic and cytokine-mediated mechanisms of resistance, recent advances underscore the critical role of cellular metabolism and metabolic crosstalk within the bone marrow niche in shaping leukemic progression and therapeutic relapse (5).

Altered metabolic programs are increasingly recognized as a hallmark of AML, contributing to its uncontrolled proliferation, impaired differentiation, and therapy resistance (6, 7, 8). A well-characterized example involves mutations in isocitrate dehydrogenase (IDH), which result in the production of the oncometabolite R-2-hydroxyglutarate from α-ketoglutarate (αKG). This metabolite inhibits αKG-dependent dioxygenases, thereby promoting leukemogenesis (9). Although targeted therapies for IDH-mutant AML are now clinically approved, targeting the metabolic vulnerabilities of AML subtypes lacking mutations in metabolic enzymes has proven difficult. For example, AML cells frequently display changes in their energy metabolism, including glycolysis, the tricarboxylic acid (TCA) cycle, and oxidative phosphorylation (OXPHOS) (7, 8, 10). Notably, OXPHOS activity is closely linked to the maintenance of leukemia stem cells, a population that drives relapse (11, 12). Unfortunately, even the newest generation of OXPHOS-targeting drugs such as IACS-010759 have shown poor clinical efficacy and significant toxicity (13). Nucleotide biosynthesis has also emerged as a key metabolic dependency in AML, with the inhibition of *de novo* pyrimidine or purine biosynthesis shown to induce differentiation and cell death in preclinical models (14, 15, 16). However, clinical translation has again been challenging, with concerns about the toxicity of existing nucleotide synthesis inhibitors (17, 18).

The limitations of existing metabolism-targeting drugs highlight the need to identify alternative strategies for selectively exploiting the metabolic vulnerabilities of AML cells. In our recent work, we have identified aspartate as one of the metabolites transferred from bone marrow stromal cells to leukemic cells, where it contributes to pyrimidine biosynthesis (19). Aspartate, a non-essential amino acid ubiquitously synthesized by mammalian cells, plays a central role not only in nucleotide synthesis, but also contributes to protein biosynthesis, redox balance, TCA cycle anaplerosis, nitrogen shuttling, and mitochondrial metabolism (20, 21). Remarkably, we found that aspartate concentrations in the bone marrow are 70- to 100-fold higher than in peripheral blood, suggesting a unique metabolic landscape supporting leukemic metabolism (19). Further supporting this, studies across multiple cancer types have implicated aspartate as a limiting metabolite under hypoxic conditions (22). Given the physiologically low oxygen tension of the bone marrow niche (23), AML cells may be particularly reliant on exogenous aspartate when growing in this environment.

Based on the established dependency of AML on pyrimidine biosynthesis and the notion that the metabolic constraints of the bone marrow microenvironment may limit aspartate synthesis, we hypothesized that targeting aspartate uptake could serve as an upstream strategy to disrupt pyrimidine metabolism more selectively and safely. Excitatory amino acid transporters (EAATs) have traditionally been studied for their role in glutamate clearance in the central nervous system, but they also transport aspartate and are increasingly recognized as key regulators of amino acid metabolism in non-neuronal tissues, including cancer (24, 25). In the current study, we investigated whether AML cells express and rely on EAATs for aspartate uptake, and whether inhibition of these transporters impacts AML cell growth or response to therapy.

## Methods

### Ethics statement

All animal procedures performed in this study were approved by the UCLouvain Institutional Animal Care and Use Committee (approval number 2021/UCL/MD/03). At the end of the experiments, mice were euthanized by CO_2_ inhalation.

Immortalized human cell lines were commercially obtained and their use for scientific research is exempt from ethical review in accordance with Belgian law. No new human material was collected in this study. Publicly available human gene and protein expression data were analyzed anonymously.

### Public datasets of gene and protein expression

We obtained values of gene expression for *SLC1A1, SLC1A2* and *SLC1A3* from BloodSpot (26) (www.bloodspot.eu, datasets: BeatAML, MILE) and Vizome (www.vizome.org, datasets: BeatAML2.0, attributes: FAB BlastMorphology, isDenovo, isTherapy). Protein levels of EAAT1, EAAT2 and EAAT3 in AML were obtained from The Cancer Genome Atlas (TCGA) (27) (proteomics.leylab.org, dataset: TMT).

### Cell lines

MA9-1 cells were obtained by retroviral infection of whole bone marrow cells with the humanized MLL-AF9 fusion oncogene as previously described (28). MA9-2 cells were obtained after crossing MLL-AF9 knock-in mice with ubiquitous GFP-expressing mice, as well as luciferase-expressing mice, and have been previously described (19). Both cell models were cultured at 37°C in a humidified incubator with ambient air and 5% CO_2_ in RPMI1640 medium (ThermoFisher, 31870074) supplemented with 10% fetal bovine serum (FBS; Sigma Aldrich, F7524), 100 I.U./mL penicillin-100 µg/mL streptomycin (ThermoFisher, 15140122), 2 mM L-glutamine (Fisher Scientific, 11510626), 20 ng/mL recombinant mouse stem cell factor, 10 ng/mL recombinant mouse interleukin 3 and 10 ng/mL recombinant mouse interleukin 6 (BioLegend Europe BV, 579708, 575708, 575508). U937, NB4, MV4-11, MOLM14, OCI-AML3, NOMO-1 and MONOMAC6 (MM6) human AML cell lines were obtained from the Leibniz Institute DSMZ and cultured at 37°C in a humidified incubator with ambient air and 5% CO_2_ in RPMI1640 medium supplemented with 10% FBS, 100 I.U./mL penicillin-100 µg/mL streptomycin and 2 mM L-glutamine. Cell line identity was validated by DNA fingerprinting and cultures were tested for mycoplasma contamination every six months using a Mycoplasma PCR Detection Kit (Abcam, ab289834). For aspartate depletion experiments, custom-made aspartate-free, glutamine-free RPMI1640 medium (ThermoFisher) was supplemented with 10% dialyzed FBS (Fisher Scientific, 11520646), 100 I.U./mL penicillin-100 µg/mL streptomycin and 2 mM L-glutamine.

### Chemicals

TFB-TBOA (Axon MedChem, 2640), UCPH-101 (Bio-Techne, 3490/50), dimethyl-aspartate (Sigma Aldrich, 456233-5G), dimethyl-alpha-ketoglutarate (Sigma Aldrich, 349631-5G), uridine (Sigma Aldrich, U3750-25G) and tyrosine (Sigma Aldrich, T8566) were added to cell cultures at the indicated concentrations. For *in vitro* chemotherapy treatment, doxorubicin (Sigma Aldrich, C6645-1G) at 30nM and cytarabine (Sigma Aldrich, D1515-10MG) at 10nM were added to cultures simultaneously.

### EAAT1 cell surface staining

Both human and mouse samples were stained with an anti-EAAT1 antibody (Alomone Labs, AGC-021) for 30 minutes at 4°C, washed with PBS (ThermoFisher, 70011044), and then incubated with a secondary antibody, anti-rabbit AF647 (Thermofisher, A-21236), for 15 minutes. Samples were analyzed by flow cytometry (FACSVerse, BD Biosciences).

### MTT assay

Cells were incubated with indicated drugs/molecules, then treated with 1.5 mg/mL Thiazolyl Blue Tetrazolium Bromide (MTT, Fisher Scientific, 15234654) for 4 hours, followed by cell lysis with 10% SDS (Sigma Aldrich, 05030-500ML-F) and overnight incubation. The absorbance was read at 560 nm with a reference wavelength of 630 nm using a Glomax (Promega). Half-maximal inhibitory concentrations (IC50) were calculated using GraphPad Prism 10.

### Cell death assay

Cells were treated with the indicated drugs or molecules for 24 hours and then harvested and stained with annexin-V-APC (BioLegend, 640941) and 7-AAD (BioLegend, 420403) according to the manufacturer’s protocol. Samples were analyzed by flow cytometry.

### Cell cycle assay

Cells were treated with indicated molecules for 6 hours, harvested and washed with PBS. The pellet was fixed with 2 mL of ice-cold 70% ethanol (Avantor, 20.821.310) in a drop-wise manner while gently vortexing, kept for 30 minutes on ice and centrifuged at 1000 x g for 5 minutes. Cells were then washed twice with PBS and incubated for 20 minutes with 200 µL of 1 µg/mL DAPI (Roche, 10236276001). Cells were washed and analyzed by flow cytometry.

### Metabolomics analysis

For metabolomics analysis by gas chromatography-coupled mass spectrometry (GC-MS), cells were treated for 6 hours with indicated molecules, harvested and washed twice in saline (NaCl 0,9%, Sigma Aldrich, 71376-1KG). The pellet was resuspended and vortexed thoroughly in 0.5 mL of methanol (kept at -20°C, Sigma Aldrich, 900688-1L), after which 0.5 mL ice-cold water and 1 mL of chloroform (Sigma Aldrich, 650471-1L) were added and samples were agitated for 15 minutes at 4°C. Cells were then centrifugated at 14,000 x g for 10 minutes at 4°C, after which the polar (top) phase was collected and dried in a SpeedVac vacuum concentrator calibrated at -100°C. Metabolites were then resuspended in 15 µL of methoxamine (Fisher scientific, 11567630) and incubated for 90 minutes at 30°C with shaking at 300 rpm. Next, 30 µL of MSTFA (Macherey-Nagel, MN701270.201) was added, samples were vortexed for 5 seconds and incubated for 30 minutes at 37°C. The mix was then centrifugated again at 14,000 x g for 10 minutes and 40 µL were transferred into a GC-MS glass vial. Samples were analyzed by GC-MS (ThermoScientific, Trace 1310, Agilent CP-Sil 8 CB-MS 30 m x 0.25 mm x 0.25 μm capillary column) on the same day of derivatization.

### Mouse model and drug treatments

8- to 10-week-old male and female C57Bl/6 mice were transplanted with 1 million murine MLL-AF9-GFP-luciferase cells (MA9-2) through intravenous injection (tail vein). Leukemia engraftment and disease progression were monitored using bioluminescence imaging on an IVIS In Vivo Imaging System (PerkinElmer) after injecting mice with D-Luciferin (ZellBio Gmbh, LUCK-1G), as previously described (19). For single agent EAAT inhibitor treatments, animals were randomized 4 days after cell injection and treated daily with UCPH-101 (20 mg/kg in 10% DMSO, 40% PEG300, 5% Tween-80, 45% saline) or TFB-TBOA (20 mg/kg in saline) through intraperitoneal injection until day 21. For combination with chemotherapy, treatment with UCPH-101 or TFB-TBOA was started at day 7 and continued for 2 weeks. Treatment with standard induction chemotherapy (iCT) started at day 9 and consisted of cytarabine (100 mg/kg in saline) given once daily for 5 days and doxorubicin (3 mg/kg in saline) given once daily for the first 3 days, via intraperitoneal injection.

### Statistical analysis

All results are reported as mean ± standard deviation (SD). Statistical significance of the difference between experimental groups was analyzed by two-tailed unpaired Student’s t-test, one-way ANOVA with Bonferroni post-hoc test or the logrank (Mantel-Cox) test for the Kaplan-Meier survival curve analyses using the GraphPad Prism 10 software. Differences were considered statistically significant at P < 0.05.

## Results

### AML cells express *SLC1A3*

Aspartate can be transported across the plasma membrane by members of the EAAT family, and particularly by EAAT1 (encoded by *SLC1A3*), EAAT2 (encoded by *SLC1A2*) and EAAT3 (encoded by *SLC1A1*) (25). We analyzed the expression of these putative aspartate transporters in human AML cells using existing databases containing gene expression data of AML patient samples, including those with different molecular subtypes. In the BEAT-AML cohort (29), we observed that *SLC1A3* is highly expressed in AML cells across molecular subtypes compared to *SLC1A2* and *SLC1A1* (**Fig. 1A, Suppl. Fig. S1A, B**). Further investigation based on the FAB classification revealed that myelomonocytic (AML-M4) and monocytic (AML-M5) AML subtypes had the highest expression of *SLC1A3* (**Fig. 1B**). We also noted that relapsed AML had higher expression of *SLC1A3* when compared to diagnostic specimens (**Fig. 1C**). Examination of the data of the MILE study (30) confirmed higher expression of *SLC1A3* compared to *SLC1A2* and *SLC1A1* in AML cells, and showed that myeloid blood cancers, including AML, express slightly higher levels of *SLC1A3* compared to lymphoid leukemias (**Suppl. Fig. S1C-E**).

**Figure 1.**
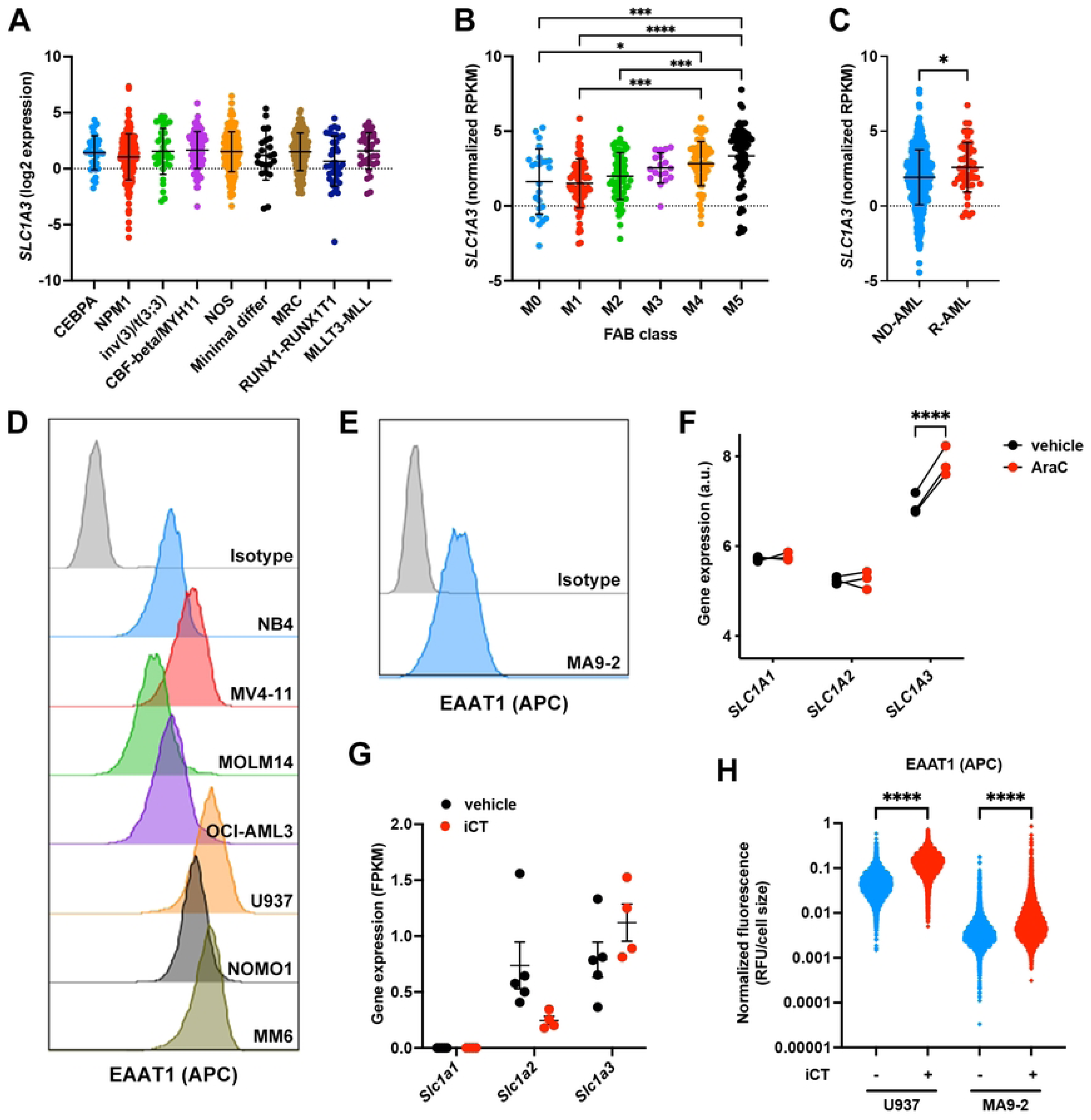
AML cells express *SLC1A3*. **A-C**. Expression of *SLC1A3* in AML cells across different patient genetic subgroups (**A**), according to FAB classification (**B**) and comparing newly diagnosed (ND) and relapsed (R) patients (**C**) in the BEAT-AML cohort. **D-E.** Histograms representing the baseline cell surface levels of EAAT1 in human AML cell lines (**D**) or a mouse AML MLL-AF9 cell line (**E**) as quantified by flow cytometry. **F.** Expression of *SLC1A1*, *SLC1A2* and *SLC1A3* by human AML patient-derived xenograft cells isolated from mice treated with cytarabine or vehicle. Data obtained from (31). **G.** Expression of *Slc1a1*, *Slc1a2* and *Slc1a3* by mouse AML cells (MA9-2 cell line) isolated from mice treated with induction chemotherapy (iCT; doxorubicin 3 mg/kg and cytarabine 100 mg/kg given in a 5+3 regimen) or vehicle. Data obtained from (19). **H.** EAAT1 cell surface protein levels in AML cells in the absence or presence of iCT drugs (10 nM cytarabine and 30 nM doxorubicin) for 24 hours, as determined by flow cytometry. Data are presented as mean ± SD. *P < 0.05; ***P < 0.001; ****P < 0.0001.

The gene expression results were confirmed at the protein level using proteomics data from The Cancer Genome Atlas (TCGA (27)), which showed the presence of EAAT1, but not EAAT2 or EAAT3, in AML patient cells, with the highest levels found in M4 and M5 subtypes (**Suppl. Fig. S1F**). Using an antibody specific for EAAT1, we confirmed the presence of this transporter on the cell membrane of different human and mouse AML cell lines. We observed that the monocytic AML cell line U937 had the highest expression compared to other AML cell lines (**Fig. 1D, E**).

To better understand the effects of therapy on *SLC1A3* expression, we re-analyzed previously published RNA sequencing datasets of human and mouse AML cells obtained from engrafted mice treated with induction chemotherapy (iCT; doxorubicin + cytarabine) or cytarabine (AraC) alone (19, 31), which showed that both human and mouse residual AML cells exhibit increased expression of *SLC1A3* after therapy (**Fig. 1F, G**). We validated this finding by treating human U937 cells and a mouse MLL-AF9-expressing AML cell line (MA9-2) with iCT *in vitro* and found that the cells that survived the chemotherapy had higher EAAT1 levels, suggesting that this transporter may play a role in chemotherapy resistance in AML (**Fig. 1H**).

### EAAT1 inhibitors reduce AML growth *in vitro*

To investigate the importance of exogenous aspartate for AML cell growth, we exposed both mouse and human cell lines to known EAAT1 inhibitors, and assessed the effects on cell health, apoptosis and proliferation. We first used increasing concentrations of TFB-TBOA (an aspartate analog and competitive inhibitor of EAAT1, EAAT2, and EAAT3 (32)) and UCPH-101 (an uncompetitive allosteric inhibitor selective for EAAT1 (33)) and measured overall cell health of MA9-1 and U937 cells using the MTT assay. Metabolic activity of both cell lines was reduced in a dose-dependent manner by the aspartate transporter inhibitors; however, UCPH-101 was found to be more potent (**Fig. 2A**). We repeated this experiment with a larger number of human and mouse AML cell lines and calculated the half-maximal inhibitory concentration (IC50) for TFB-TBOA and UCPH-101. All tested cell lines responded similarly to both molecules, with UCPH-101 demonstrating consistently higher potency (**Fig. 2B**).

**Figure 2:**
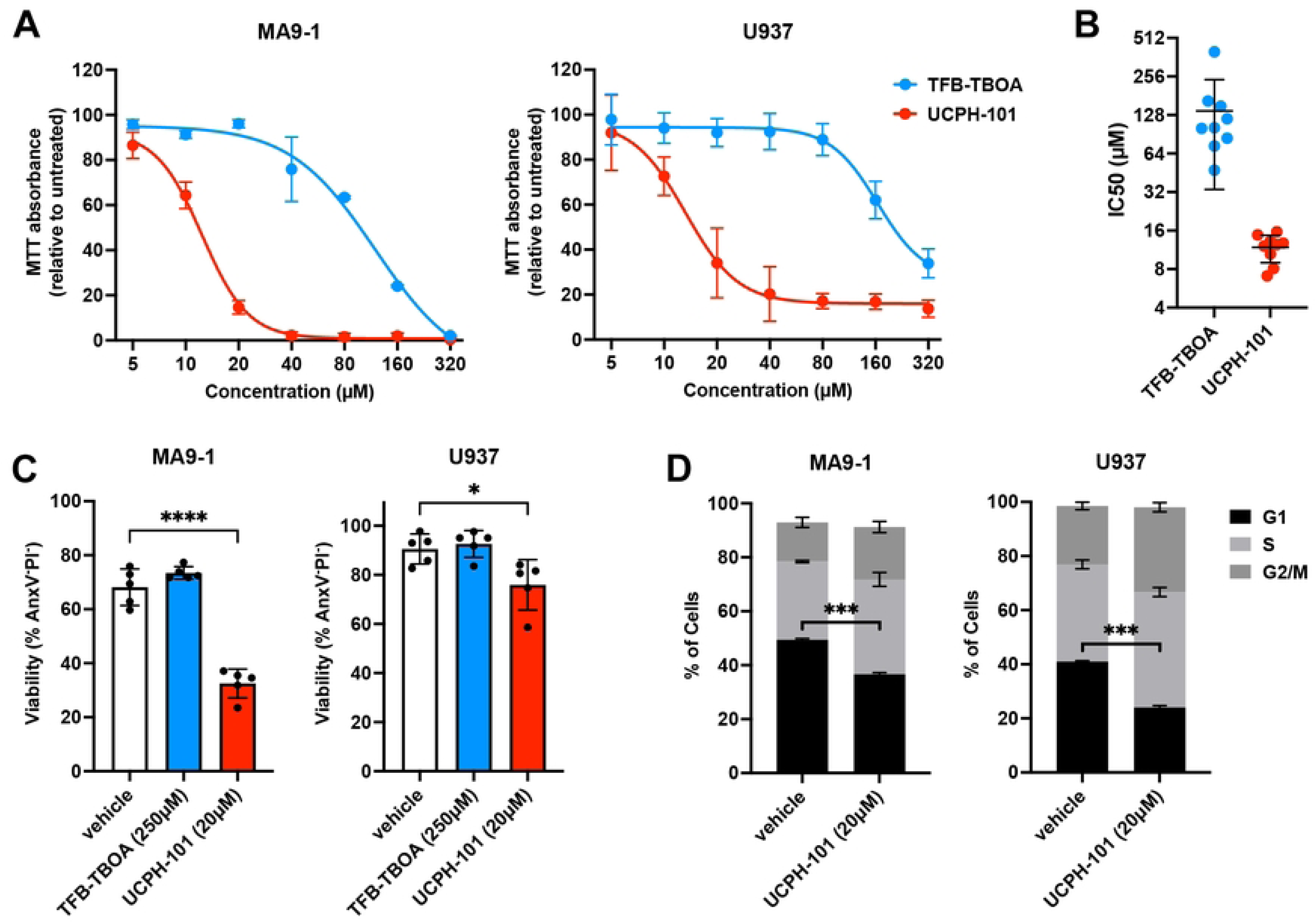
EAAT1 inhibitors reduce AML growth *in vitro*. **A.** Overall health of AML cells exposed to increasing concentrations of TFB-TBOA or UCPH-101 for 24 hours, as determined by the MTT assay. **B.** Half-maximal inhibitory concentration (IC50) for TFB-TBOA and UCPH-101 across 9 different human and mouse AML cell lines, based on the MTT assay. Each dot represents a different AML cell line. **C.** Viability of AML cells treated with TFB-TBOA (250 µM) or UCPH-101 (20 µM) for 24 hours as measured by flow cytometry. **D.** Cell cycle analysis of AML cells treated with UCPH-101 (20 µM) for 6 hours as measured by flow cytometry. Data are presented as mean ± SD. *P < 0.05; ***P < 0.001; ****P < 0.0001.

Given that the MTT assay measures overall metabolic health, we further explored how the inhibitors affected AML cells by measuring apoptosis and performing cell cycle analysis. In both the mouse MA9-1 and human U937 cell line we observed increased apoptosis when cells were exposed to 20 µM of UCPH-101 for 24 hours, whereas 250 µM of TFB-TBOA did not affect cell viability (**Fig. 2C**). Cell cycle analysis, conducted after 6 hours of UCPH-101 treatment (20 µM) to prevent cell death, revealed fewer cells in the G1 phase and a corresponding increase in cells in the S and G2/M phases, indicating a dysregulation of the cell cycle (**Fig. 2D**). Taken together, these results show that AML cells are sensitive to aspartate transporter inhibitors, with UCPH-101 showing the highest potency.

### UCPH-101 kills AML cells in an aspartate-independent manner

To evaluate the metabolic effects of UCPH-101, we treated MA9-1, MA9-2, and U937 cell lines with this inhibitor and quantified intracellular metabolites by GC-MS. After 6 hours of incubation, aspartate levels were reduced, along with tyrosine and several TCA cycle metabolites. In contrast, essential amino acid levels were increased, demonstrating a significant metabolic effect of the inhibitor (**Fig. 3A, B**).

**Figure 3:**
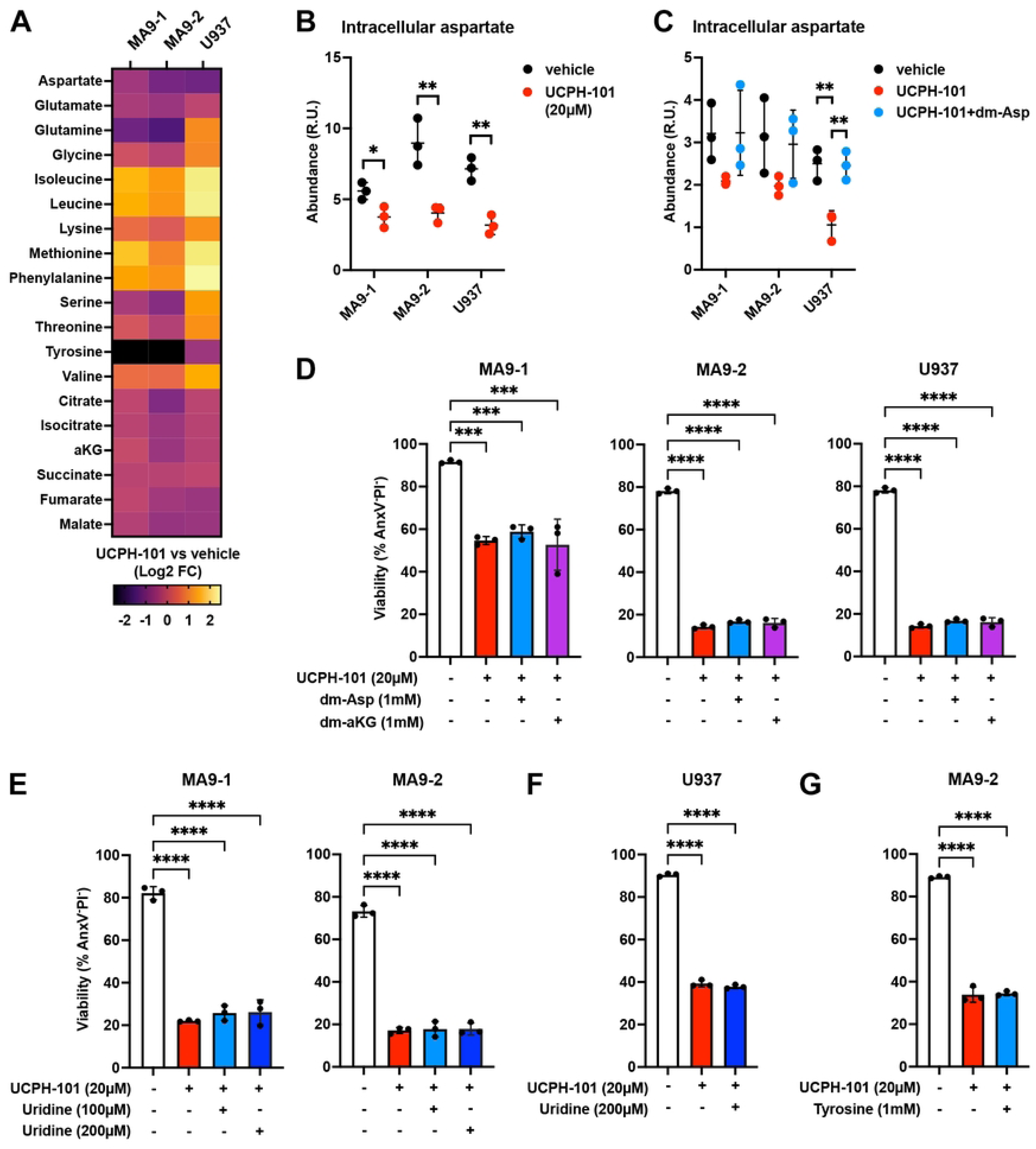
UCPH-101 kills AML cells in an aspartate-independent manner. **A.** Heatmap showing relative intracellular levels of different metabolites in AML cells treated with UCPH-101 (20 µM) for 6 hours as measured by GC-MS. aKG: alpha-ketoglutarate. **B.** Quantification of intracellular levels of aspartate in AML cells treated with UCPH-101 (20 µM) for 6 hours, as measured by GC-MS. R.U.: Relative Units. **C.** Quantification of intracellular levels of aspartate in AML cells treated with UCPH-101 (20 µM) with or without dimethyl-aspartate (dm-Asp; 1 mM) for 6 hours, as measured by GC-MS. **D.** Viability of AML cells treated with UCPH-101 with or without dimethyl-aspartate (dm-Asp) or dimethyl-alpha-ketoglutarate (dm-aKG) at the indicated concentrations for 24 hours, as measured by flow cytometry. **E-F.** Viability of mouse MLL-AF9 AML cells (**E**) or human U937 cells (**F**) treated with UCPH-101 with or without uridine at the indicated concentrations for 24 hours, as measured by flow cytometry. **D.** Viability of mouse MLL-AF9 AML cells treated with UCPH-101 with or without tyrosine at the indicated concentrations for 24 hours, as measured by flow cytometry. Data are presented as mean ± SD. **P < 0.01; ***P < 0.001; ****P < 0.0001.

We hypothesized that blockade of aspartate uptake was responsible for the observed metabolic changes. To counteract the effects of the inhibitor, we treated the AML cell lines with both UCPH-101 and dimethyl-aspartate (dm-Asp), a membrane-permeable form of aspartate. GC-MS analysis confirmed rescue of intracellular aspartate levels by dm-Asp in mouse and human AML cell lines treated with UCPH-101 (**Fig. 3C**). However, addition of dm-Asp did not rescue cell viability, suggesting that UCPH-101 induces AML cell death through a mechanism unrelated to intracellular aspartate levels (**Fig. 3D-F**). Based on the other metabolic effects observed after treating AML cells with UCPH-101 (**Fig. 3A**), we supplemented the medium with dimethyl-alpha-ketoglutarate (dm-aKG), uridine, or tyrosine in the presence of UCPH-101. We found that none of these supplements restored cell viability (**Fig. 3D-J**). These data show that when used at 20 µM, UCPH-101 kills AML cells through a mechanism independent of aspartate uptake, suggesting that either EAAT1 has other substrates, or UCPH-101 has other targets.

### AML cells do not require extracellular aspartate *in vitro*

Due to the potential off-target effects of UCPH-101, we lowered the concentration to 10 µM, which did not show cytotoxicity when used as a single agent (**Fig. 4A**). Given that electron transport chain activity is essential for *de novo* aspartate synthesis (34, 35) and hypoxia has been shown to limit the ability of cancer cells to synthetize aspartate (22), we tested whether lowering oxygen tension would increase sensitivity to EAAT1 inhibition. MA9-2 mouse AML cells, when cultured at 21%, 2% or 0.5% oxygen, did not exhibit differences in baseline viability, but at lower oxygen tensions the cells became slightly more sensitive to UCPH-101 (**Fig. 4A**).

**Figure 4:**
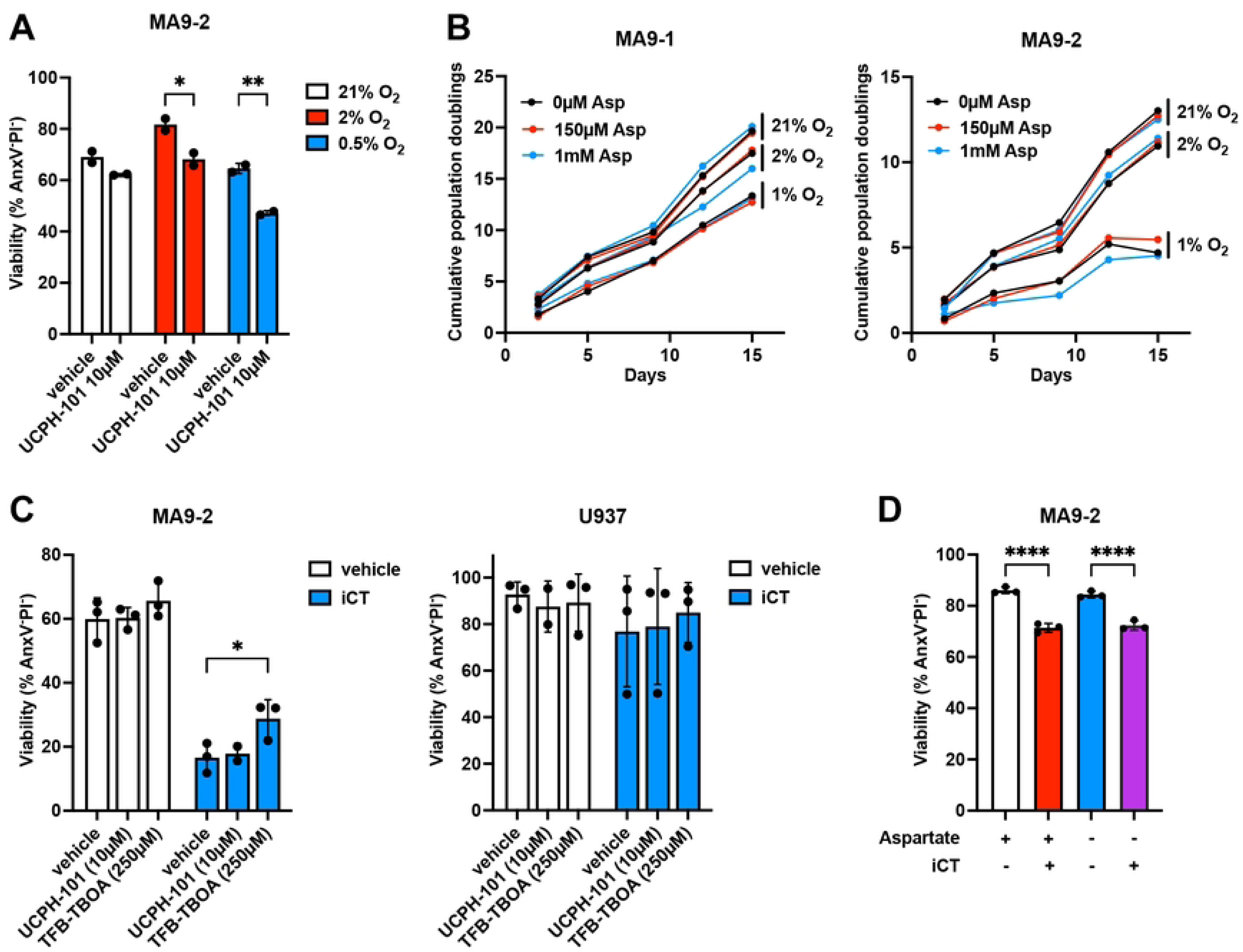
AML cells do not require extracellular aspartate *in vitro*. **A.** Viability of mouse MLL-AF9 AML cells cultured for 24 hours at 21%, 2% or 0.5% oxygen tension in the absence or presence of UCPH-101 (10 µM), as measured by flow cytometry. **B.** Cumulative population doublings of mouse MLL-AF9 AML cells cultured at 21%, 2% or 1% oxygen tension in the absence or presence of aspartate in the culture medium at the indicated concentrations. **C.** Viability of AML cells treated with vehicle, TFB-TBOA (250 µM) or UCPH-101 (10 µM) in the absence or presence of induction chemotherapy (iCT) drugs (10 nM cytarabine and 30 nM doxorubicin) for 24 hours, as measured by flow cytometry. **D.** Viability of AML cells treated with vehicle or iCT drugs in the absence or presence (150 µM) of aspartate in the culture medium for 24 hours, as measured by flow cytometry. Data are presented as mean ± SD. *P < 0.05; **P < 0.01; ****P < 0.0001.

To better understand the importance of aspartate as a nutrient for AML cells, we next cultured MA9-1 and MA9-2 mouse AML cell lines in regular RPMI-1640 medium (containing 150 µM aspartate), aspartate-free medium, and medium containing 1 mM aspartate (to mimic aspartate levels in the bone marrow plasma (19)). Each of the cultures was further maintained at different oxygen levels (21%, 2%, 1% O_2_). Population doubling analysis revealed that AML cells grew normally under all conditions, regardless of extracellular aspartate levels (**Fig. 4B**). Expectedly, lower oxygen availability decreased cell growth, but it did not increase sensitivity to extracellular aspartate depletion. These findings support the notion that UCPH-101 impacts AML cell growth and viability through its off-target effects and show that AML cells do not require exogenous aspartate for *in vitro* growth.

Since exposure to iCT increased EAAT1 levels in AML cells (**Fig. 1H**), we exposed MA9-2 mouse AML cells to iCT in combination with the EAAT1 inhibitors or with aspartate depletion from the medium. However, neither UCPH-101 at 10 µM, TFB-TBOA at 250 µM nor aspartate-free medium sensitized AML cells to iCT (**Fig. 4C, D**), indicating that exogenous aspartate is also not required for AML cells to survive the consequences of chemotherapy *in vitro*.

### Inhibition of EAAT1 does not alter AML growth or chemotherapy response *in vivo*

While AML cells do not appear to require extracellular aspartate when cultured *in vitro*, we hypothesized that in the bone marrow microenvironment these cells may depend on aspartate as a nutrient for their growth or survival following iCT treatment. Since direct depletion of aspartate in the bone marrow plasma would be challenging, we tested whether treatment of mice with TFB-TBOA or UCPH-101 could impact aspartate uptake by cells in the bone marrow.

To this end, we treated control C57Bl/6 mice daily with TFB-TBOA (for 4 or 8 days) or UCPH-101 (for 2 days) and isolated the bone marrow plasma. Analysis by GC-MS revealed elevated levels of aspartate in the bone marrow plasma after treatment of mice with the EAAT1 inhibitors, suggesting an accumulation of the metabolite due to reduced cellular consumption and thus validating the effect of the compounds *in vivo* (**Fig. 5A, B**). In contrast, when we measured the levels in the peripheral blood, no changes were observed (**Fig. 5A**).

**Figure 5:**
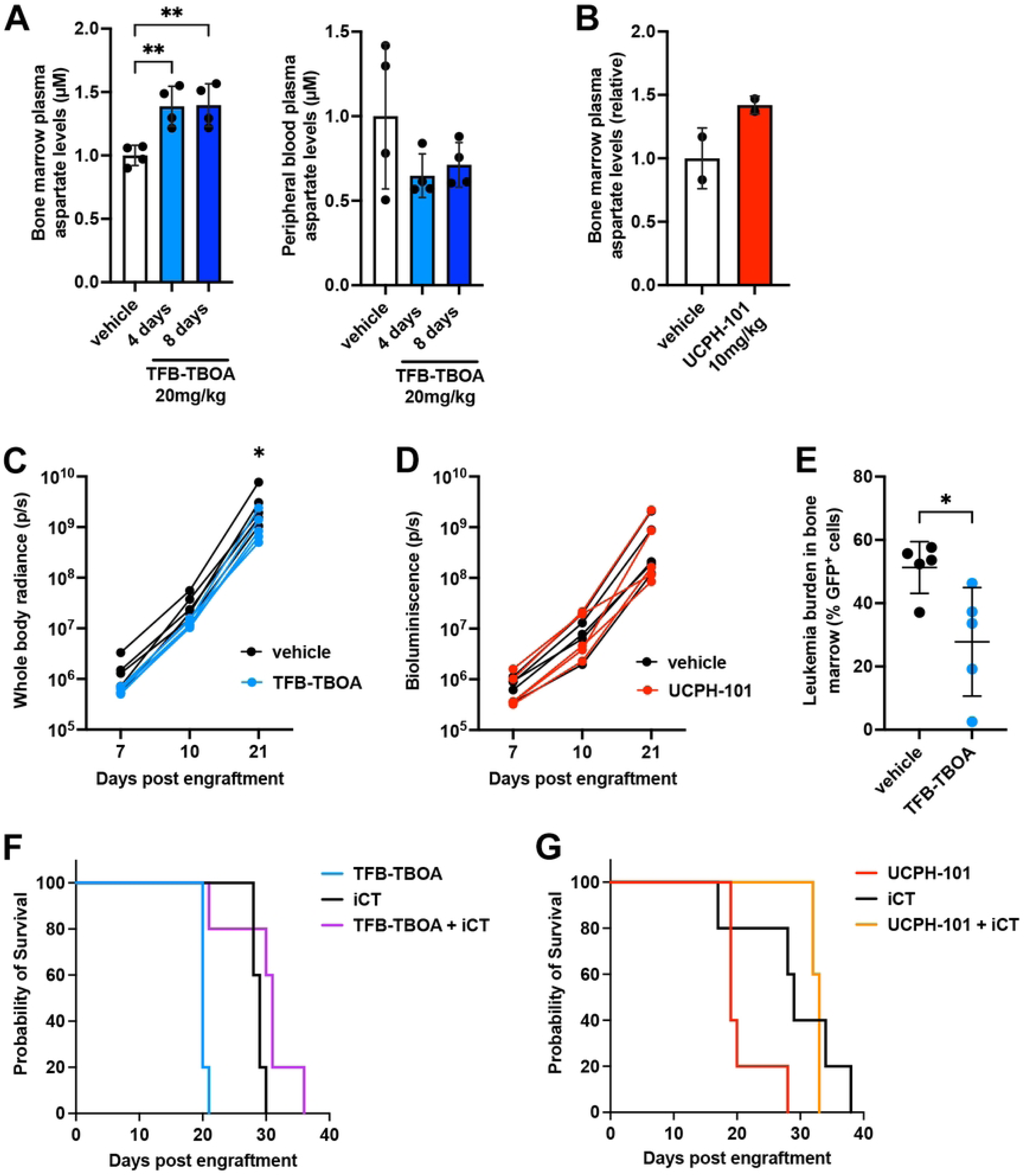
Inhibition of EAAT1 does affect AML cells *in vivo*. **A.** Relative aspartate levels in the bone marrow plasma or peripheral blood plasma of mice treated with vehicle for 8 days or 20 mg/kg TFB-TBOA for 4 or 8 days, as measured by GC-MS. **B.** Relative aspartate levels in the bone marrow plasma of mice treated with vehicle or 10 mg/kg UCPH-101 for 2 days, as measured by GC-MS. **C-D.** Leukemic burden as measured by whole body radiance of mice engrafted with MA9-2 GFP- and luciferase-expressing AML cells and treated daily with vehicle, 20 mg/kg TFB-TBOA (**C**) or 10 mg/kg UCPH-101 (**D**) starting at day 4 post engraftment. **E.** Leukemic burden in the bone marrow at day 10, as measured by flow cytometry, of mice engrafted with MA9-2 GFP- and luciferase-expressing AML cells and treated daily with vehicle or 20 mg/kg TFB-TBOA starting at day 4 post engraftment. **F-G.** Kaplan-Meier survival curves of mice engrafted with MA9-2 AML cells and treated daily with vehicle, 20 mg/kg TFB-TBOA (**F**) or 10 mg/kg UCPH-101 (**G**) starting at day 7 post engraftment, with or without concomitant induction chemotherapy (iCT; doxorubicin 3 mg/kg and cytarabine 100 mg/kg given in a 5+3 regimen) treatment started at day 9 post engraftment. Data are presented as mean ± SD. *P < 0.05; **P < 0.01.

We then engrafted non-irradiated C57Bl/6 recipient mice with MA9-2 GFP- and luciferase-expressing AML cells and treated them with TFB-TBOA, UCPH-101 or their respective vehicle starting at day 4 post-engraftment. Tracking of leukemic burden by bioluminescence imaging demonstrated that AML cells were able to grow normally in the presence of either of the aspartate transporter inhibitors, although TFB-TBOA did appear to slightly reduce AML cell growth (**Fig. 5C, D**). To further investigate this, we sacrificed mice at day 10, after 6 days of TFB-TBOA treatment, and analyzed the number of GFP^+^ AML cells in the bone marrow, which confirmed a modest but significant reduction in leukemic burden (**Fig. 5E**).

Next, we combined treatment of AML-engrafted mice with EAAT1 inhibitors (starting at day 7) and iCT (doxorubicin and cytarabine given in a 5+3 regimen, starting at day 9). While iCT prolonged mouse survival, neither TFB-TBOA nor UCPH-101 further enhanced this response (**Fig. 5F, G**), thus showing that inhibition of aspartate transporters does not impact the response of AML cells to chemotherapy *in vivo*.

## Discussion

In the current study, we used small molecule inhibitors, metabolic profiling, functional analyses and *in vitro* and *in vivo* AML models to demonstrate that EAAT aspartate/glutamate transporters are not a critical metabolic dependency in AML. Despite its expression in AML cells and upregulation with chemotherapy treatment, EAAT1 inhibition fails to significantly impair leukemic cell viability or enhance the efficacy of standard chemotherapy.

Based on our previous observations that the bone marrow plasma contains an elevated concentration of aspartate and that residual AML cells persisting after iCT highly depend on pyrimidine synthesis (19), which requires aspartate as a substrate, we hypothesized that AML cells rely on aspartate/glutamate transporters of the EAAT family for their growth and survival. We found that AML cells across genetic backgrounds express EAAT1, with the highest expression seen in M4 and M5 AML subtypes, but detected only very low levels of the other EAATs. *In vitro*, pharmacological inhibition of EAAT1 with UCPH-101 reduced intracellular aspartate levels and AML cell viability, but only when the inhibitor was used at 20 µM or higher. In contrast, concentrations of 1-1.5 µM have been shown to fully block aspartate or glutamate uptake in other cell models including HEK293 cells and astrocytes (36, 37, 38). In line with this, we found that the cell death induced by 20 µM UCPH-101 was aspartate-independent and thus likely the consequence of off-target effects of the inhibitor.

Given that low oxygen tensions limit the ability of cancer cells to synthetize aspartate (22), exogenous aspartate may become more important as a nutrient under hypoxic conditions such as those found in the bone marrow (23). However, our findings suggest that even under hypoxia, AML cells exhibit a high degree of resilience to aspartate deprivation. Similarly, we found that the inhibition of EAATs had only mild effects on AML growth or response to chemotherapy *in vivo*. These findings underscore the intrinsic metabolic plasticity of leukemic blasts, which, like their normal hematopoietic counterparts, are adept at surviving and proliferating in the oxygen-poor bone marrow microenvironment. Moreover, a recent study indicates that cancer cells rapidly adapt to a reduction in intracellular aspartate levels by establishing a new steady-state and that the negative effects of electron transport chain inhibition only manifest when aspartate levels fall below a critical threshold (39). In addition to EAATs, AML cells may use other transporters to take up aspartate, and could additionally replenish their aspartate pools not only by synthesizing aspartate via glutamate-oxaloacetate transaminases, but also by converting asparagine to aspartate, recycling aspartate from intracellular protein turnover, importing extracellular proteins via macropinocytosis or engaging in metabolic exchange through cell-cell interactions (20). Such metabolic plasticity would allow AML cells to circumvent blockade of a single pathway, further illustrating the challenge of targeting individual metabolic nodes in malignancy.

The lack of effect of EAAT1 inhibition in AML stands in contrast to a previous report which identified this transporter as a metabolic vulnerability in T-cell acute lymphoblastic leukemia (40). However, this study implicated EAAT1 as a mitochondrial glutamate/aspartate transporter involved in glutamine-to-aspartate conversion rather than as a cell surface transporter involved in aspartate uptake, and used UCPH-101 at 25 µM without validating whether its effects were due to aspartate deprivation. A direct comparison of their results to ours is therefore not evident.

Taken together, our findings suggest that while extracellular aspartate and its transport via EAAT1 may contribute to aspartate pools in AML, this transporter does not constitute a targetable vulnerability for this cancer type. Future therapeutic strategies may need to involve the simultaneous disruption of several aspartate acquisition routes to overcome metabolic redundancy. However, such approaches must carefully consider toxicity to normal hematopoietic progenitors, which may share these adaptive traits. Continued efforts to identify selective vulnerabilities in AML metabolism, possibly through synthetic lethality or metabolic bottlenecks unique to malignant cells, will be essential for advancing more effective therapies.

## Authorship contributions

H.A.T. and N.v.G. conceptualized the study and designed experiments. H.A.T., N.B., F.L. and L.L. performed experiments. H.A.T., N.B. and N.v.G. analyzed and interpreted results. H.A.T. and N.v.G. wrote the manuscript text and prepared the figures. N.v.G. acquired funding and supervised the research. All authors reviewed the manuscript.

## Acknowledgements

We thank Isabelle Gerin, Francesco Caligiore and Guido Bommer from the Metabolic Research Group at de Duve Institute (UCLouvain, Belgium) for access to and help with GC-MS analysis. We also thank Nicolas Dauguet from the CYTF platform at the de Duve Institute for help with flow cytometry analysis, and all members of the Cellular Metabolism and Microenvironment Laboratory for helpful discussions.

## Funding

This work was supported by the Belgian Foundation Against Cancer [F/2020/1440, F/2024/2556], the Fund for Scientific Research - FNRS [F.R.S.-FNRS; A5/5-CQ/135, M4/1/2/5-MIS/BEJ] and the de Duve Institute. H.A.T. was supported by a doctoral research fellowship from the Fund for Research Training in Industry and Agriculture (FRIA) [1.E.027.22] and by a Bourse du Patrimoine from the UCLouvain. L.L. was supported by a FRIA doctoral research fellowship [1.E017.24], N.B. was supported by a post-doctoral *Chargée de Recherches* fellowship from the F.R.S.-FNRS [1.B.027.24].

## Notes

### Competing Interest Statement

The authors have declared no competing interest.

